# Correction for spurious taxonomic assignments of k-mer classifiers in low microbial biomass samples using shuffled sequences

**DOI:** 10.1101/2025.06.18.660363

**Authors:** Shan Sun, Anthony A. Fodor

**Affiliations:** Department of Bioinformatics and Genomics, College of Computing and Informatics, UNC Charlotte, Charlotte, NC, USA

## Abstract

**Background:** With the increased use of shotgun metagenome and metatranscriptome sequences in characterizing the microbiome, accurate taxonomic classification of sequencing reads is essential for interpreting microbial community composition and revealing differential microbial signature between groups. K-mer based classifiers such as Kraken2 provide high speed and sensitivity, and are commonly used for low microbial biomass samples. However, their performance can be compromised by specific sources of error without proper parameter settings and incorporation of controls.

**Methods:** In this study, we analyzed six sequencing datasets of human tumor biopsies with Kraken2 and investigated how shared compact hash codes (i.e., identical hash codes across different k-mers), hash collision and the structure of reference databases can contribute to false positive taxonomic assignments in low biomass samples.

**Results:** We demonstrated that in samples with high non-microbial DNA noise, the classified taxa of Kraken2 in sequencing reads are significantly correlated with that of shuffled sequences using the default setting. These taxa showed a similar distribution as those overrepresented in the hash table construction of the reference database. Incorporation of controls using shuffled reads can separate significant taxa with more robust differences from those more affected by background noise. Although the confidence thresholds needed to minimize noise varied with taxa, a minimum value of 0.2 can also help reduce misclassifications.

**Conclusion:** Our findings highlighted the need for caution when interpreting low-abundance or unexpected taxa in sequencing datasets of low microbial biomass samples. This work contributes to a more comprehensive understanding of the limitations of k-mer based classification tools and provides practical guidance for improving accuracy in microbiome research.

## Introduction

The number of studies investigating the roles of the human microbiota in cancer has increased rapidly in recent years. However, the accurate characterization of microbiota in cancer and normal tissues has been hindered by the contamination of human genomes. Compared to the low microbial biomass in tissues, the large amount of human genome DNA can cause false positive detection of microbes, which has led to controversial findings of microbial signatures associated with cancer types, placenta and blood (1–3). Many studies have discussed the importance of the extensive removal of human genome sequences and curation of a cleaner database in minimizing spurious findings (2, 4). Besides these efforts and a strict contamination control in sample collection, we explored how optimization of classification protocols can also contribute to increasing the confidence of taxonomic classification.

The taxonomy classification methods of sequencing reads generally belong to the following three categories: k-mer based methods (e.g., Kraken) (5), marker-gene based methods (e.g., MetaPhlan) (6) and alignment based methods (e.g., BLAST) (7). Alignment based methods are rarely used for large-scale metagenomics and metatranscriptomics data due to their low speed.

Marker-gene based methods rely on the finding of a set of marker genes and generally yield low classification rates in low microbial biomass samples. As a result, k-mer based methods are more commonly used in low microbial biomass samples such as studies for the human cancer microbiome.

K-mer based methods can achieve high sensitivity in detecting microorganisms in low biomass samples. But they need to be utilized with due caution and the results need to be interpreted carefully. One of the main drawbacks of k-mer based methods is false classification because of hash collision. K-mer based methods load a hash table of the hash codes of k-mers and corresponding taxonomy ID into RAM in order to achieve rapid classification. To alleviate the burden of required RAM, a compact hash table is usually used. For example, in the standard Kraken2 database, 32 bits of key-value pairs are used where the hash codes of k-mers were used as keys (15 bits) and the taxonomy IDs were used as values (17 bits) (5). Using the default setting of Kraken2, where detection of one 35-kmer theoretically leads to the classification of the sequencing read, the false positive rates are therefore expected to be high. While Kraken2 offers the option of confidence score cutoff (the number of detected k-mers / the number of total k-mers in the read), it has not been a default setting and is not always used in running pipelines.

This is especially problematic in complex or low-biomass samples, where the signal-to-noise ratio is low, and spurious assignments can obscure meaningful biological patterns. For example, in the study of human microbiome, pre-processing steps usually include alignment and removal of non-microbial DNA such as human genome DNA. However, due to the genetic diversity of human genomes, regional variation and incomplete parts, it is difficult to ensure that all non-microbial DNA were depleted from tissue samples that are highly enriched in human DNA (4). This is even more challenging in environmental samples that lack clear definitions of references for non-microbial DNA.

Previous studies that evaluated the performance of microbiome classification tools mostly used simulated reads and mock microbial communities (8–10), without particular analysis of their performance in low microbial biomass samples and the mechanisms underlying spurious false positive assignments. In this study, we selected six publicly available human tumor datasets that were not constrained to controlled access, including sequencing of biopsy samples of brain tumor, lung cancer, ovarian cancer, colorectal cancer, colon cancer and mucoepidermoid carcinoma. We evaluated how the settings and database can lead to incorrect findings of microbial signatures and how choices of parameters and re-evaluation of findings can generate findings with higher confidence. By characterizing the conditions under which spurious classifications arise, we aim to provide insights into the limitations of current methods and databases, offer recommendations for improving classification accuracy of microbiome in low biomass samples, and contribute to the growing effort to enhance the robustness and reproducibility of microbiome analyses.

## Methods

We selected six publicly available sequencing datasets of human tumor tissues (Table 1). The sequencing reads were analyzed with Kraken2 with the standard database (Dec 2024) with both default settings and settings with a range of cutoffs for confidence score. To build controls with similar nucleotide compositions, we shuffled the order of nucleotides of each sequencing read without changing the read length or GC contents across all the samples. Besides the shuffling at the nucleotide level (1-mer), we also tested the performance by shuffling k-mers with k ranging from 2 to 35 in one sample. We then used Kraken2 for taxonomic classification of the shuffled reads. The human genomes were not removed from these reads in order to estimate the influence of host genome contamination. The statistical analyses were performed using R (version 4.3.1). The PCoA and PERMANOVA analyses were generated using R package ‘vegan’. The taxonomic differences between one cancer type and the rest were analyzed with Wilcoxon tests. The associations between taxa and the number of minimizers were calculated using modified Kraken2 scripts to generate text files instead of binary files for the databases. Scripts used in this study are available at GitHub (https://github.com/ssun6/Error_Control_Cancer_Microbiome).

**Table 1.** Information of the sequencing datasets analyzed.

| BioProject | Sample type | Source | # Reads (average) | Read length | Sample size |
| --- | --- | --- | --- | --- | --- |
| PRJNA988034 | RNA-Seq | Colorectal tumor | 14,921,154 | 150 | 34 |
| PRJNA1009135 | RNA-Seq/WXS | Brain tumor | 59,484,862 | 300 | 10 |
| PRJNA1014965 | RNA-Seq | Mucoepidermoid carcinoma | 55,036,219 | 200 | 20 |
| PRJNA1168164 | RNA-Seq | Ovarian tumor | 44,879,997 | 300 | 35 |
| PRJNA1186843 | RNA-Seq | Lung cancer | 152,015,602 | 300 | 18 |
| PRJNA1204136 | RNA-Seq | Colon tumor | 42,096,393 | 300 | 30 |

## Results

### Random reads are more likely classified to certain bacteria due to database bias

In the standard Kraken2 database, 32 bit key-value pairs are used to store the taxonomy and compact hash codes, with 17 bit for NCBI taxonomy ID and 15 bit for the hash code of 35-mer of sequencing reads. To generate the compact hash code using the default setting, a canonical 31- mer minimizer is calculated first, 7 nucleotides are masked from the 31-mer and Murmurhash3 function is used to generate the value and the first 15 bits of the hash code are used as keys for the database. This compact hash table allowed a much higher processing speed and reduced RAM use compared to Kraken1 that used 96-bit key-value pairs. However, because of the limited bits for compact hash codes, there is an increasing chance of misclassification of random k-mers that were not present in the genomes (Figure S1).

Because of the known genome database bias towards bacteria associated with health and industrial use, we hypothesized that random k-mers are more likely to be classified as those bacteria. To identify the bacteria prone to high misclassification rates, we counted the number of minimizers for each NCBI taxa ID and found that the numbers were not evenly distributed across taxa (Figure 1). Some taxa had higher than average numbers, for example, *Streptomyces* comprised 2.43% of total minimizers and that is 759 times of the average percentage (5.3E-3%). *Pseudomonas* comprised 1.38% of total minimizers, and Bacillus comprised 0.30%. Genera *Bradyrhizobium* and *Rhizobium* that includes many nitrogen-fixing rhizosphere species comprised 0.27% and 0.16% respectively.

**Figure 1.**
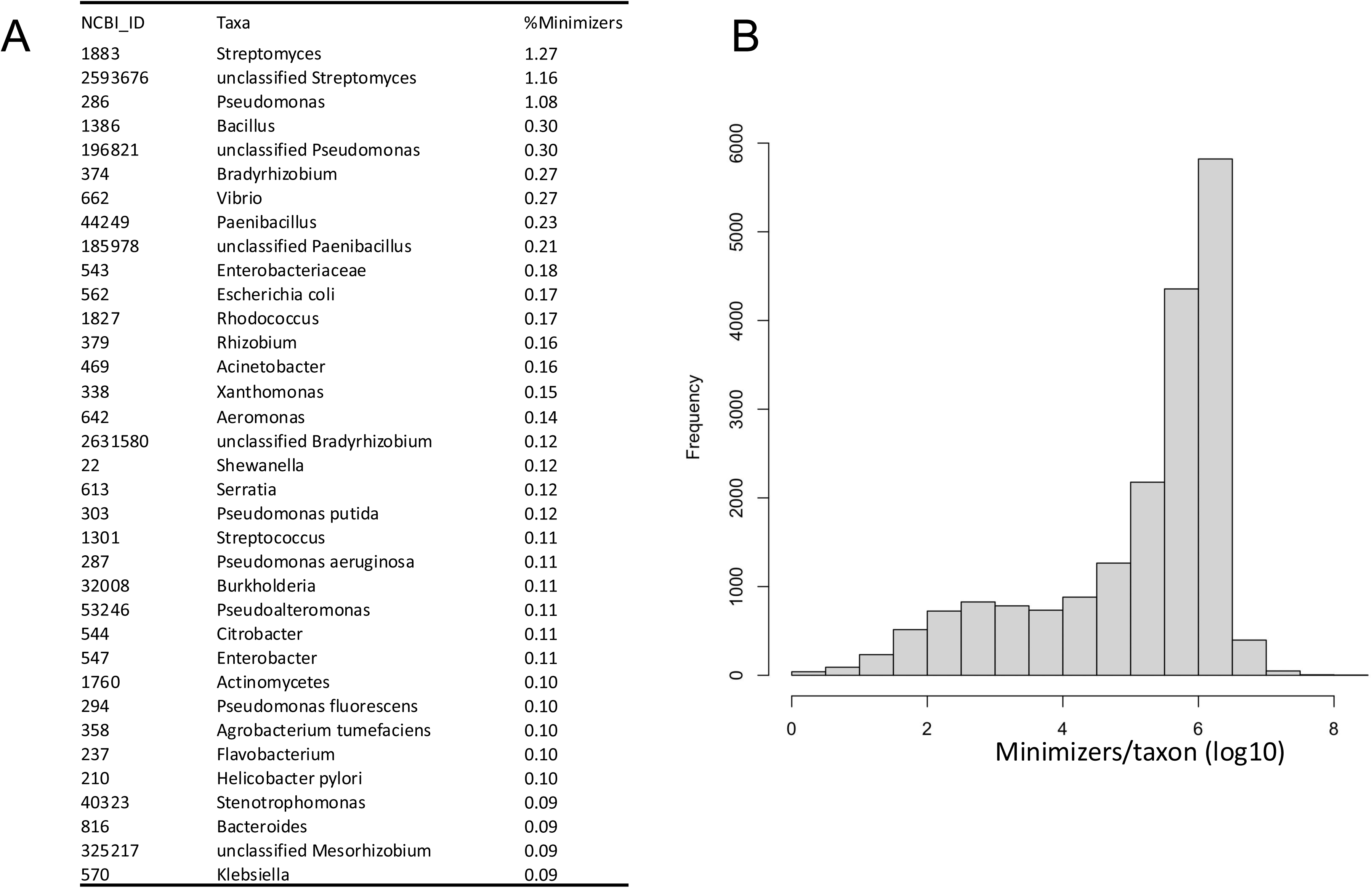
Top taxa that are overrepresented in the hash table of the standard Kraken2 database (Dec 2024) (A) and the overall distribution of hash values assigned to each taxon (B).

### Construction of negative controls using shuffled sequences

To construct negative controls that preserve local sequence composition while disrupting biological signals, we generated randomized reads by shuffling k-mers within each original sequencing read. First, we tested the performance of randomizing k-mers with a range of k from 1 to 35 using sample SRR13502479. Shuffling with a 1-mer completely scrambles the sequence and, if the sequence was in a one-dimensional array, is the equivalent of shuffling the array.

Higher k values break the sequence prior to shuffling. With a k value of 2, the sequence would be expressed as a series of 2-mers and those 2-mers would then be shuffled. We compared the taxonomic classification of shuffled and unshuffled sequencing reads at the genus and species level, and observed discrete jumps in the correlation coefficients at k values that are multiples of 3 (e.g., 3, 6, 9, etc.) (Figure 2). This periodic pattern likely reflects the underlying structure of coding sequences in bacteria genomes, where codons constitute the fundamental units of translation. Shuffling k-mers at lengths aligned with codon boundaries may preserve more biological information and have higher classification rates than non-multiples of 3. These findings underscore the importance of careful k-mer selection in benchmarking studies. Negative controls by shuffling k-mers with a k as non-multiples of 3 will disrupt the reading frame more and potentially better represent random noise. Shuffling with a k as multiples of 3 will create controls retaining some biological information, which can be used for more aggressive controls for overclassification.

**Figure 2.**
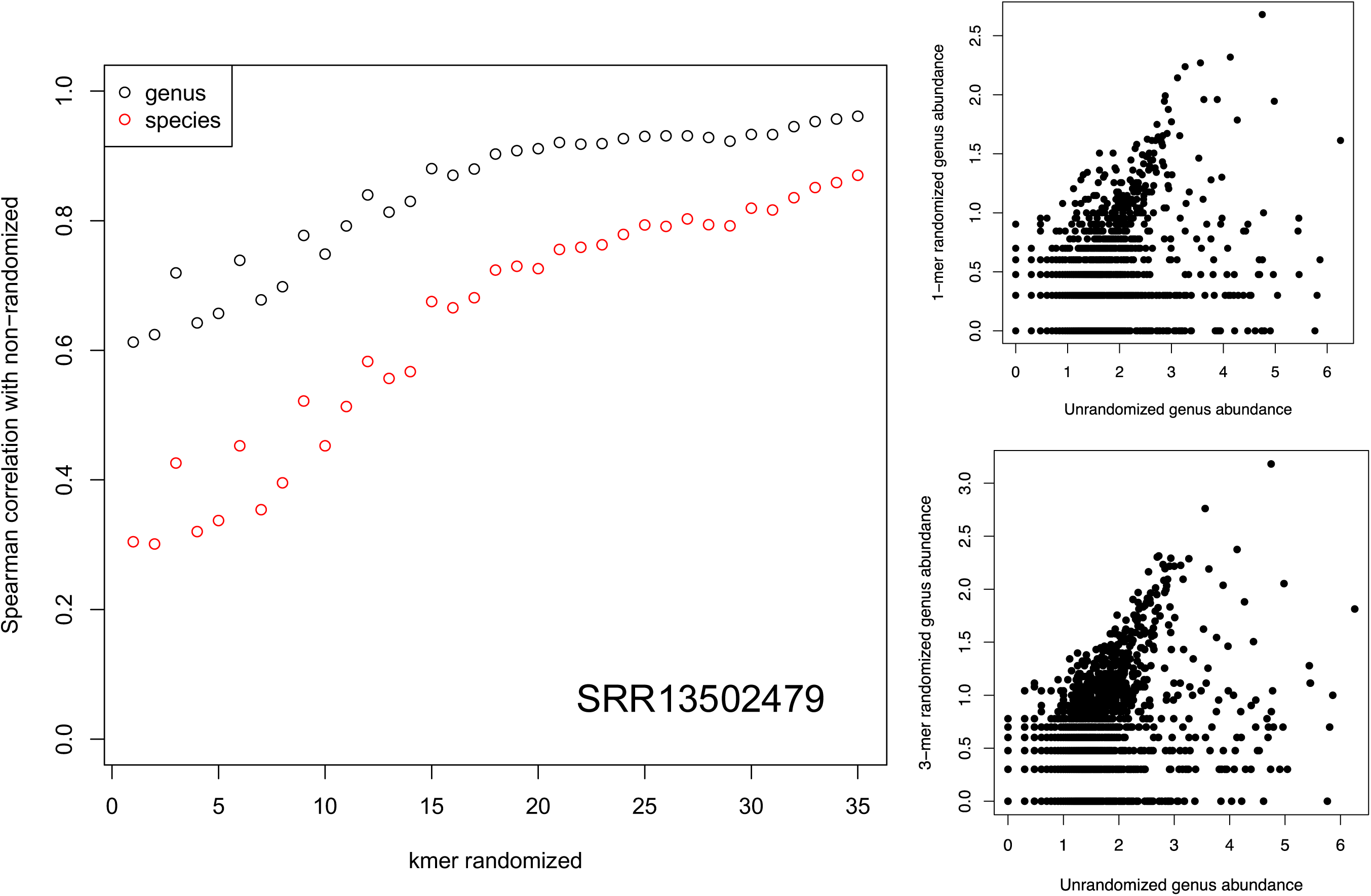
The correlation between taxonomic compositions of unrandomized reads and reads randomized by shuffling the order of k-mers with k from 1 to 35. (A) Spearman’s rho of unrandomized reads and reads randomized with k from 1 to 35. (B) Scatterplots of genus abundance of unrandomized reads and 1-mer and 3-mer randomized reads.

### The microbial taxonomic profiles of human tumor reads are similar to randomized reads

To estimate potential overclassification rates, we collected six publicly available datasets that sequenced human tumor tissue samples that are low in microbial biomass and enriched in human genome DNA, including brain tumor, lung cancer, ovarian cancer, colorectal cancer and mucoepidermoid carcinoma (Table 1). Their sequencing depth are shown in Figure 3C. We used Kraken2 with the default settings to characterize the microbial profiles of these samples. On average, 1.5% reads were classified as bacteria. The numbers of total classified reads and reads classified as bacteria are shown in Figure S2. We observed that the genus level compositions were clustered by tumor type with PCoA (Figure 3A). PERMANOVA tests suggested that the taxa composition was significantly associated with tumor types with a R^2^ of 0.383 and a P-value of 0.001. To verify if this clustering was from actual differential microbial signature or an artifact of nucleotide composition variation by tumor types, we shuffled the order of nucleotides (1-mer) in each read to create randomized reads as controls. We found that the microbial compositions of randomized reads remained clustered by cancer types and the PERMANOVA tests remained significant (R^2^ = 0.624, P-value = 0.001) (Figure 3B). This indicates that the nucleotide composition itself could explain a large fraction of the microbial variation by tumor type. We then analyzed the GC contents of the sequencing reads from different human tumor tissues (Figure 3D), and found that the GC contents were significantly different across tumor types (ANOVA, P <2e-16), which may have contributed to the clustering especially in the randomized reads.

**Figure 3.**
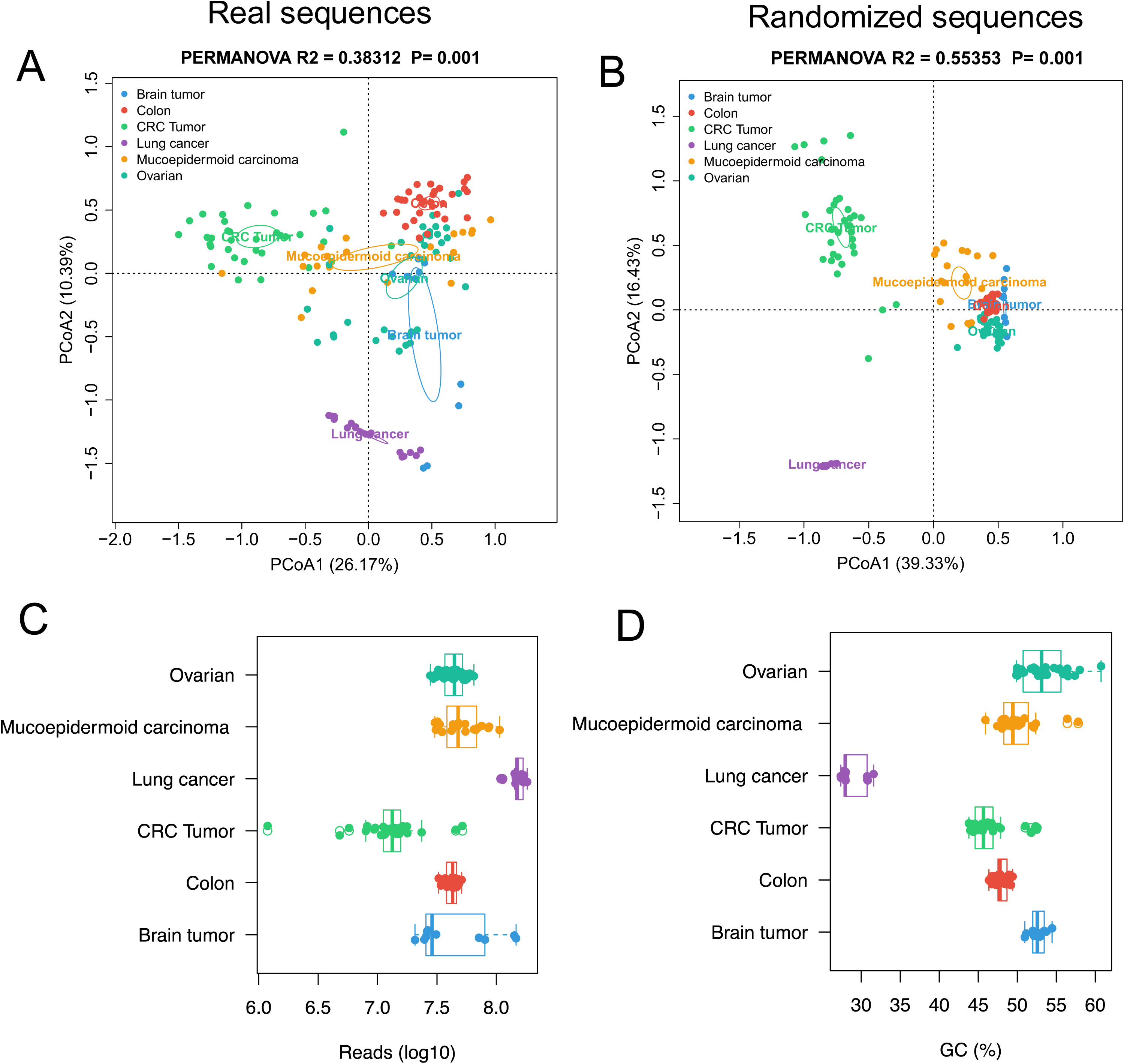
PCoA of genus composition of six cancer datasets. (A) Real reads analyzed with Kraken2 with the default setting. (B) Randomized reads analyzed with Kraken2 with the default setting. (C) Sequencing depth and (D) GC contents by datasets.

We also compared the individual genus abundance of randomized sequencing reads to that of the unrandomized reads in each tumor type and found that they were significantly correlated in all six tumor types (Spearman’s correlation, FDR<0.1). The average correlation coefficients are around 0.5 for brain tumor, colorectal cancer, mucoepidermoid carcinoma and ovarian tumor, while lung cancer has the highest correlation coefficient of 0.7 (Figure 4A). In the examples (Figure 4B) of one randomly picked sample per tumor type, many high-abundance genera had similar abundance in randomized and unrandomized samples, indicating that their abundance estimation from the unrandomized sequencing samples were likely flawed and exaggerated. The taxa of similar high abundance in randomized and unrandomized samples included Streptomyces, Pseudomonas, Bacillus and Mycobacterium (Figure 4). Rhizobium and Bradyrhizobium were also among the genera displaying similar trends. Rhizobium and Bradyrhizobium are genera of soil bacteria with many species involved in nitrogen fixation and they are therefore very unlikely human tissue symbionts.

**Figure 4.**
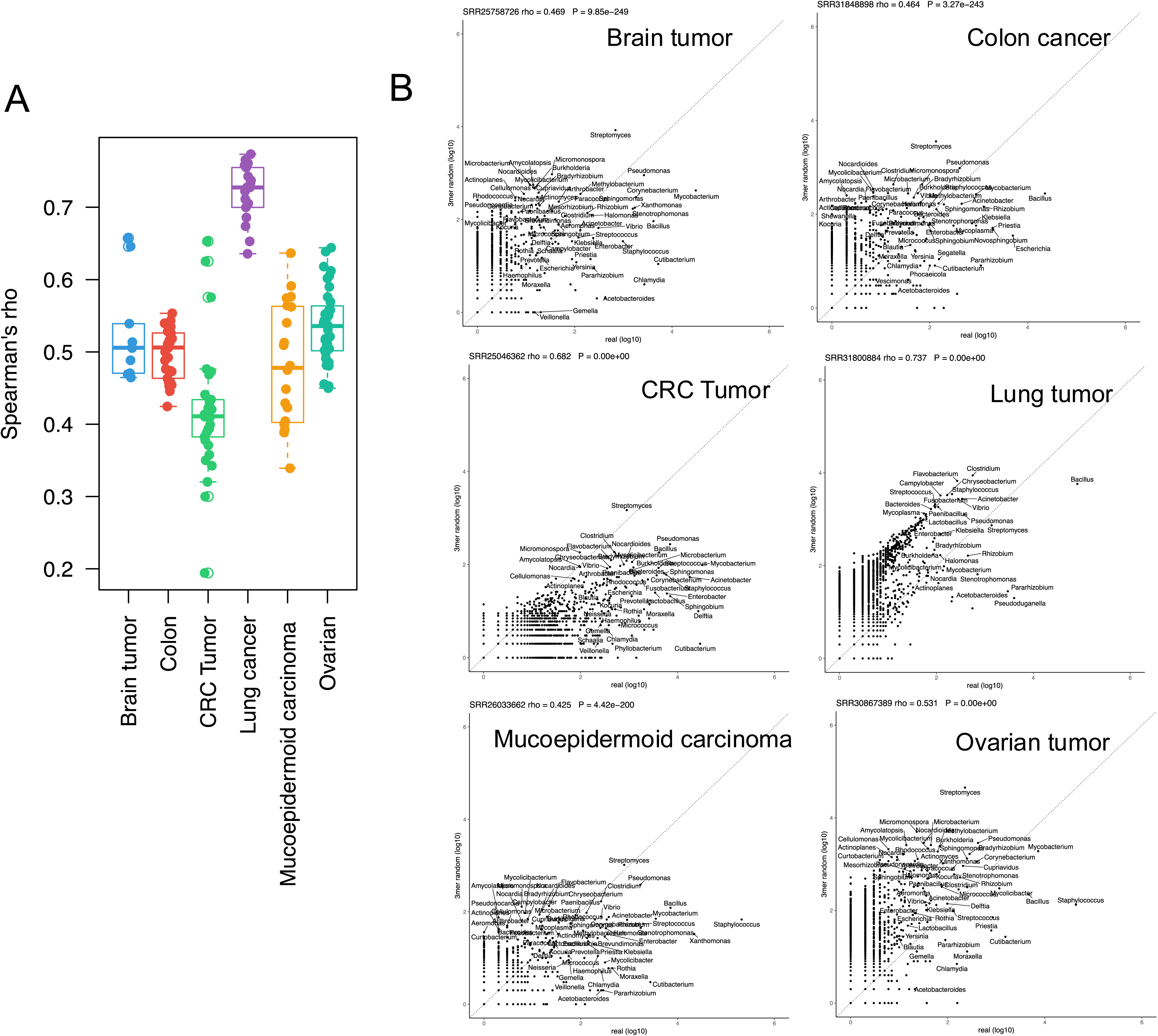
Genus composition of randomized reads were significantly correlated with unrandomized reads.

To test if the unbalanced numbers of minimizers across different taxa contributed to the over- classification of some microorganisms, we calculated the correlations between the abundance of genera and the number of their minimizers in the database in both unrandomized and randomized samples. In general, for both raw and randomized reads, there were non-trivial correlations between the relative abundance and the number of minimizers of each genus in the database across all cancer types (Figure 5). This suggests that the bias of reference database and the hash collision rate together may have led a background noise in the classification of the unrandomized reads similar to those in the randomized reads.

**Figure 5.**
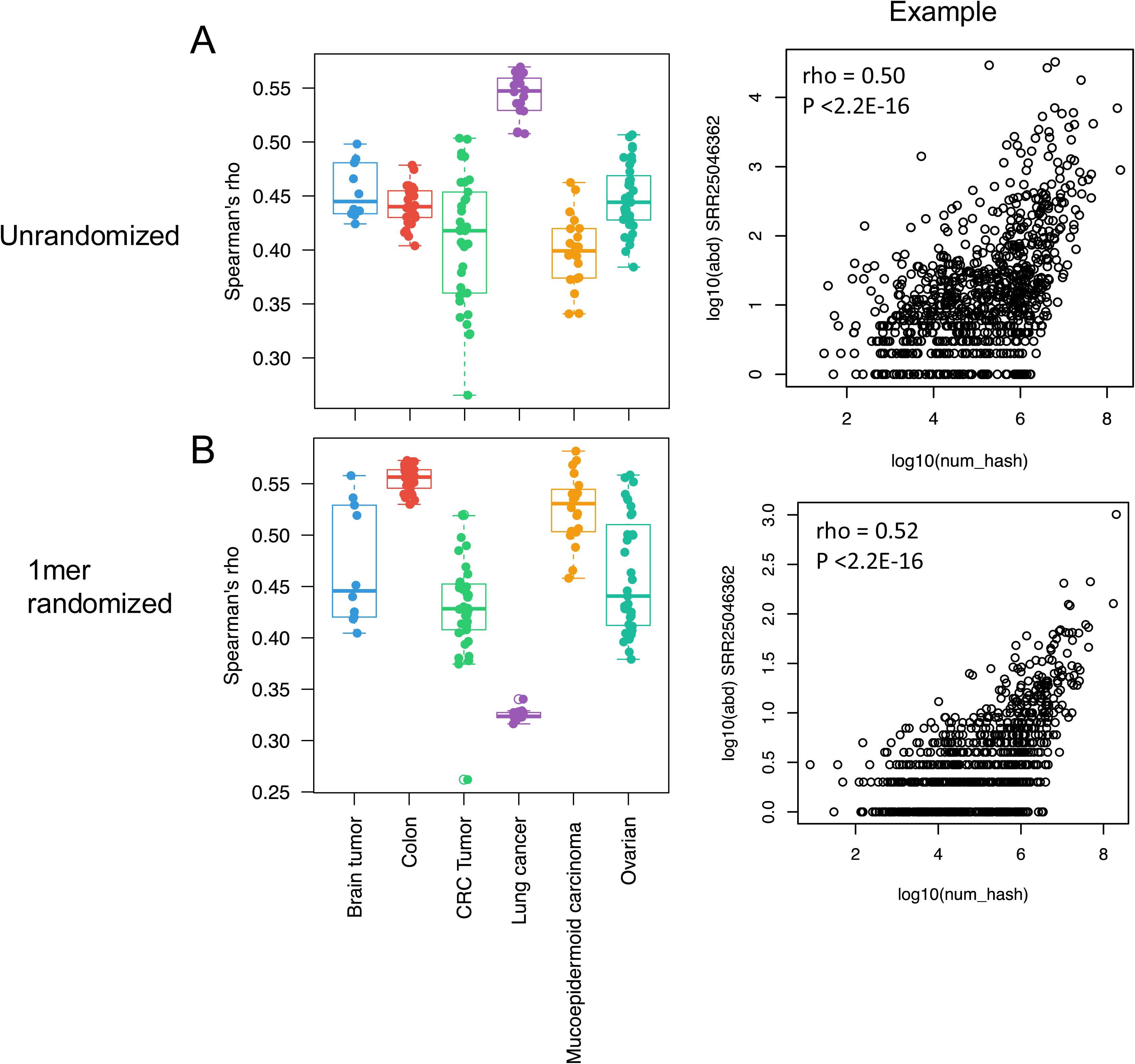
The correlation of genus abundance using (A) unrandomized reads and (B) 1-mer randomized reads in human biopsy samples and the hash abundance of genus in the Kraken2 standard database.

### Inferences of unrandomized reads were subject to systematic errors in low microbial biomass samples

To evaluate if the similar taxonomic profiles in randomized and unrandomized reads will lead to similar inferences in statistical results, we next used the Wilcoxon test to determine the taxa that can differentiate one tumor type from the rest. We found a large number of significant differential taxa for tumor types in both unrandomized and randomized reads (FDR < 0.1) (Table 2). We compared the directional log10 P-values of taxa between unrandomized and randomized reads and found the taxa separating one tumor type from the others in the unrandomized data are generally different from those in the randomized data with a few exceptions including *Streptomyces*, *Limnobacter* and *Staphylococcus* (Figure S3).

**Table 2.** The number of significant taxa between one cancer type and the others in real and randomized datasets.

|  | Real data | Randomized reads data |
| --- | --- | --- |
| Brain tumor | 190 | 1130 |
| Colon tumor | 811 | 1167 |
| Colorectal tumor | 596 | 2737 |
| Lung cancer | 957 | 3250 |
| Mucoepidermoid carcinoma | 570 | 761 |
| Ovarian tumor | 619 | 2403 |

\*P-values were from Wilcoxon tests of one cancer type and the test. P-values were adjusted with the Benjamini-Hochberg method. A cutoff of 0.1 was used for determining significance.

In an attempt to remove these systemic artifacts, we subtracted the taxa abundance of the randomized reads from the abundance of the unrandomized reads, and then performed statistical analyses on the subtracted abundance analyzing the difference between one cancer type and the rest. We compared the differential taxa estimated using subtracted abundance to those estimated by unrandomized abundance, and found that the correlation varied across cancer types. The colorectal tumor datasets had stronger correlations with Spearman’s rhos of 0.606 and 0.514 (Figure 6), followed by mucoepidermoid carcinoma (rho = 0.27), lung tumor (rho = 0.259) and ovarian tumor (rho = 0.17), while brain tumor had the weakest correlations (rho = 0.02) (Figure 6). This suggests that the inferences are more robust in colorectal tumor samples with more microbial biomass compared to other samples.

**Figure 6.**
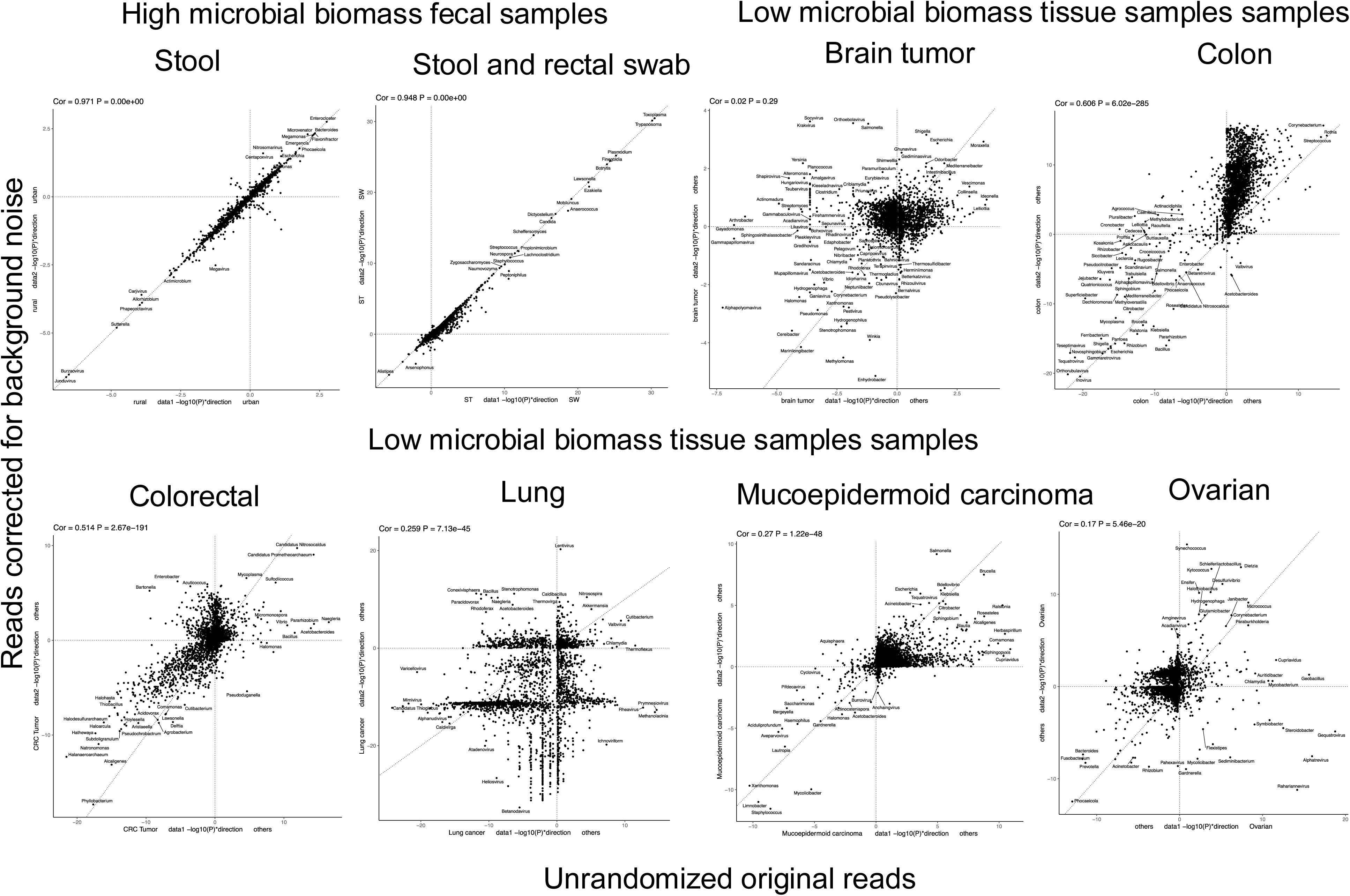
Comparison of statistical analyses results in original raw reads and reads corrected for background noise. Statistical analyses were performed to analyze the difference between one tumor type and the rest. X and Y axes are –log10(P-value) multiplied by +1/-1 to specify the direction of changes.

To compare these results to high microbial biomass samples, we ran a similar set of inference comparison analyses between subtracted reads and unrandomized reads using human stool and rectal swab samples in two publicly available datasets (Fig. 6, top left panels) (11, 12). In the first study, we compared the differential taxa separating rural to urban resident using subtracted reads or unrandomized reads, and found the results are highly similar (rho = 0.971). In the second study analyzing the differences between stool and rectal swab microbiome, a strong correlation (rho = 0.948) was observed as well.

Compared to these strong correlations observed in high microbial biomass samples, the findings in low microbial biomass samples showed poor consistency between the inferences made with subtracted and unrandomized reads. This suggests that in low microbial biomass samples, the inference made with classification of sequencing reads without control for systems errors were less reliable and the findings need careful post hoc examinations. This analysis also provides a way to examine the robustness of microbial changes. The colon and colorectal datasets had better consistency than other datasets, likely because the biopsies from these locations had more microbial biomass compared to other locations. The taxa on the identity line (remained significant after background subtraction) in the colon dataset included virus and *Escherichia* that are known to be associated with colon cancer (13–15), demonstrating the utility of this method.

### The change of classification rates across a range of confidence scores

The default settings of Kraken2 assign classification to a read with at least one hash of k-mer found in the database. With the large amount of shared compact hash codes and hash collision across k-mers, this likely leads to many misclassified reads especially in samples with a high fraction of non-microbial DNA reads such as human genome reads. There is a function in Kraken2 that can set a confidence score for the percentage of k-mers classified to one taxon before assigning classification to the read. For example, one 150 bp read has 116 35-mers, and a confidence score of 0.5 requires 58 35-mers to be classified to the same taxon before labeling the read as from that taxon.

To explore how the number of classified reads changes with different cutoffs, we ran Kraken2 on both unrandomized and randomized sequencing reads from the six datasets with different cutoffs (Figure 7). For genus Streptomyces that comprises most minimizers, a cutoff of 0.4 was needed to have zero classification in the randomized reads across all cancer types in this study. This cutoff varied slightly across different cancer types, with 0.15 for lung cancer, 0.2 for colon cancer, 0.25 for brain tumor, ovarian tumor and mucoepidermoid carcinoma, and 0.4 for colorectal cancer. *Pseudomonas* and *Bacillus* required 0.3 across all six cancer datasets, and *Bradyrhizobium* required 0.25. They also had lower cutoffs for randomized lung cancer sequences and higher cutoffs for colorectal cancer datasets. The cutoff variation may be related to the differences in the number of sequencing reads per sample and the different nucleotide compositions across tissue types. The lung cancer data had the lowest cutoff but the highest number of sequencing reads. It is also important to note that the number of classified reads in unrandomized samples decreased with an increase of confidence score cutoff, especially with Bacillus (Figure 7). This indicates that a higher than necessary cutoff could reduce the sensitivity of detecting certain taxa.

**Figure 7.**
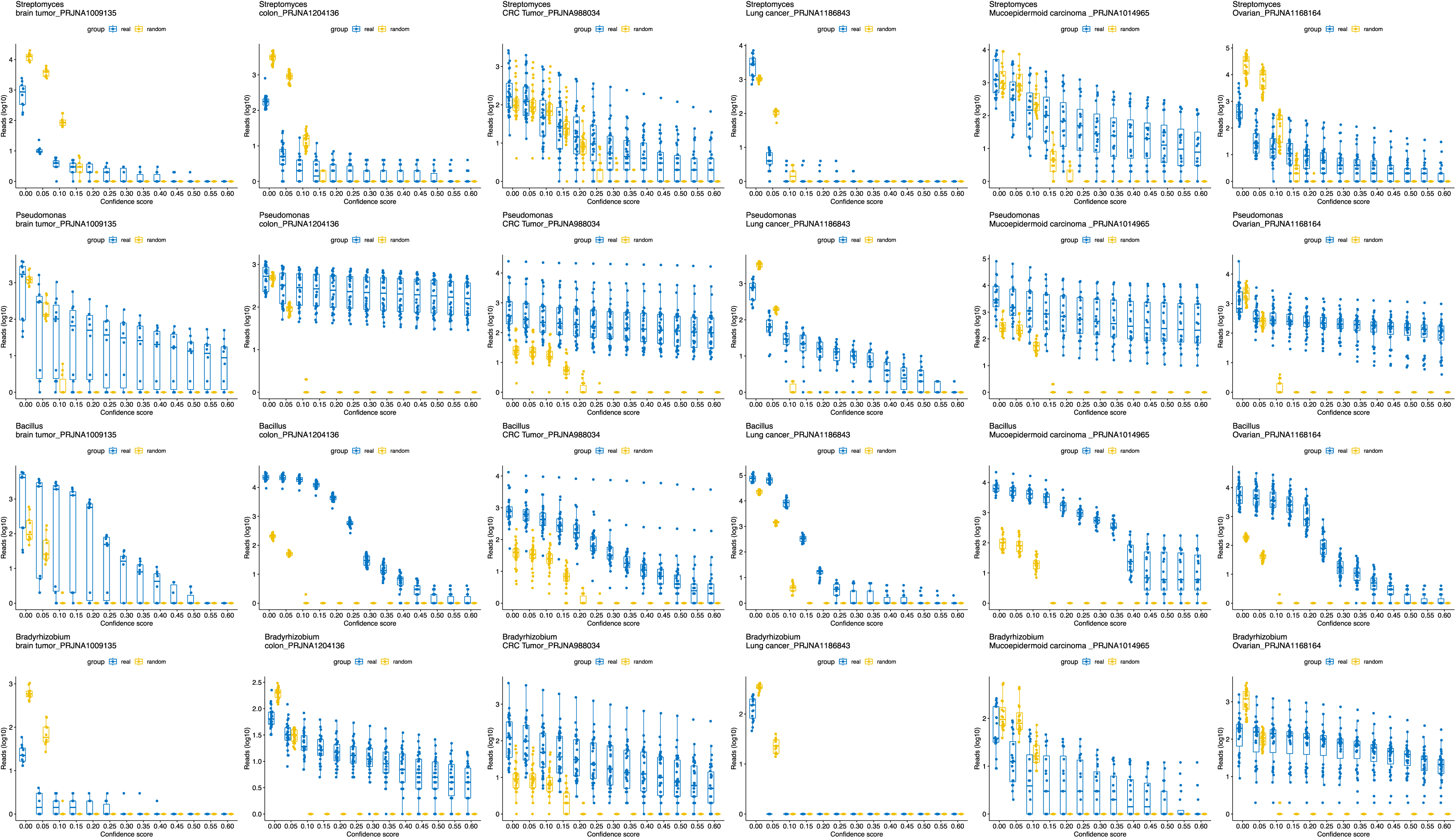
The changes of the number of reads classified to the overrepresented taxa in both real and randomized sequencing reads in each cancer dataset across different cutoffs for the confidence score.

## Discussion

In this study, we utilized Kraken2 to characterize the potential microbiome existing in the tumor tissues of six publicly available sequencing datasets and their controls built by randomizing the order of nucleotides in each sequence. We found that the characterized taxonomic profiles were clustered by cancer types, similar to previous findings that there are unique microbial signatures specific to major cancer types in a retracted study (1). However, we observed clustering by cancer type using shuffled reads as well, indicating that the clustering was likely an artifact of different nucleotide compositions instead of real microbial signatures. The consistency between the microbial taxonomic profiles of unrandomized reads and randomized reads further supported the existence of systemic classification errors.

One of the primary sources of error may have stemmed from limitations in the construction of reference databases and the trade-off between pipeline efficiency and accuracy. Classifications tools based on k-mer profiles are computationally efficient and sensitive in detecting microbes and thus are more commonly used for low microbial biomass samples compared to other types of classification tools. However, to reduce the RAM needed to store hash codes of k-mers, a compact hash table is often constructed for the reference database, which usually have a much smaller set of unique compact hash codes compared to the number of unique k-mers.

Another important contributor to misclassification is the uneven representation of taxa in reference databases. Some species, particularly clinically or industrially relevant bacteria, are represented by a much larger number of genomes compared to less-studied taxa. In the construction of hash table database, more genomes and higher k-mer diversity possibly leads to a larger fraction of hash codes assigned to those taxa such as *Streptomyces*, *Pseudomonas* and *Bacillus* in this study. This imbalance creates a bias: reads from non-microbial DNA (e.g., human genome) and environmental contamination are more likely to be misclassified to these microbes compared to others. This was confirmed by the high abundance of these taxa in the randomized reads. The number of k-mer hashes in a reference database can be conceptually viewed as a Bayesian prior distribution, randomized sequences without biological information tend to match taxa in proportion to their representation in the database (Fig. 1). Modifying the database to a more uniform hash distribution could reduce the over-classification of taxa represented more in existing datasets. Approaches including reweighting hashes or retraining classification models could also make taxonomic classifications less dependent on their distribution in the database.

Our analyses suggested a few approaches to be incorporated into microbiome analysis of low biomass samples to reduce systemic errors. First, randomized reads can be used as negative controls to distinguish true biological signal from background noise. These reads contain no valid biological information and should, in theory, be unclassifiable. Classification of these randomized reads to specific taxa indicates a systematic tendency toward overclassification, especially if the reads classified are in a similar magnitude as the real sequences.

For statistical analyses of low biomass samples, the inferences generated using real reads with randomized background subtracted can be used to distinguish biologically relevant signals from the likely background noise that needs further verification. In our statistical tests using colorectal cancer datasets, *Escherichia coli*, a taxon known to be associated with colorectal cancer, yields consistent inference from both the raw and subtracted reads. This observation suggests a potential strategy for identifying differential taxa in low biomass samples: focusing on taxa that produce similar signals before and after background subtraction. If the inferences generated using real reads are substantially different from those generated using real reads with randomized background removed, the inferences should be further verified.

Second, a confidence score cutoff should be used for classification of low microbial biomass samples with k-mer based pipelines. As previously discussed, due to the diversity of human genomes, it is difficult to ensure that all host DNA were aligned and removed before classifying microbiome in human tissues (4). It is likely a similar case for other samples with low microbial biomass but high non-microbial DNA. Based on our analysis, a cutoff of 0.2 is recommended for general classification of microbial taxonomic composition. This finding is consistent with previous analyses using simulated microbial communities to estimate classification errors (16, 17). To achieve a higher accuracy for specific microbial species such as Streptomyces, a cutoff of 0.4 is recommended.

Finally, filtering based on read counts and relative abundance before any normalization can be helpful in minimizing false positives. If a lower or no cutoff is needed to increase sensitivity for detecting certain low-abundance species, the findings need to be traced back to sequencing read to eliminate the cause of hash collision before publishing the results. The findings of low abundance taxa and unexpected taxa (e.g., nitrogen-fixing rhizosphere taxa in human tissues) should be examined and interpreted with caution. For example, *Bradyrhizobium* has been identified as a bacterial contaminant in DNA extraction kits and reagents (18).

While our findings highlighted the potential mechanism of systemic errors in k-mer classification pipelines and suggested approaches to reduce errors, several limitations should be noted. First, we note that shuffling sequences is an extreme form of background correction that disrupts biologically meaningful patterns observed in real taxa, and 1-mer shuffling may not be the most appropriate null model, which may explain the lack of concordance of inferences in Fig. S3. The question remains what makes the most appropriate null model: 1-mer shuffles, 3-mer shuffles, or alternative bootstrapping techniques that better preserve patterns in microbial sequences. This question warrants further investigation. Nonetheless, the persistence of clustering and certain taxa even in 1-mer shuffled sequences provides a valuable indication of artifactual patterns in microbial classification. Second, the true taxonomic compositions of the studied samples were unknown, which complicates the validation of inferred misclassifications and makes it difficult to calculate the real false positives and false negatives. Third, the GC contents could have contributed to the clustering by cancer type, but the underlying mechanisms of enrichment of different taxa in different cancer types remained undetermined. Finally, the analysis in this study does not specifically detect and remove contaminating microorganisms introduced during sample processing, which need to be removed before downstream analyses as well.

## Conclusion

In this study, we demonstrated that the large number of shared k-mer hash codes and database composition can significantly influence microbial taxonomic classification outcomes in low microbial biomass samples. Because of the challenges in filtering external DNA from microbiome DNA, the remaining contamination reads can be classified into spurious species genomes that are enriched in reference database and with similar nucleotide composition. These findings underscore the importance of using negative controls such as randomized reads to assess misclassification rates and incorporating confidence thresholds into classification pipelines in order to increase classification robustness and establish accurate and reproducible interpretations.

---

**Figure S1.**
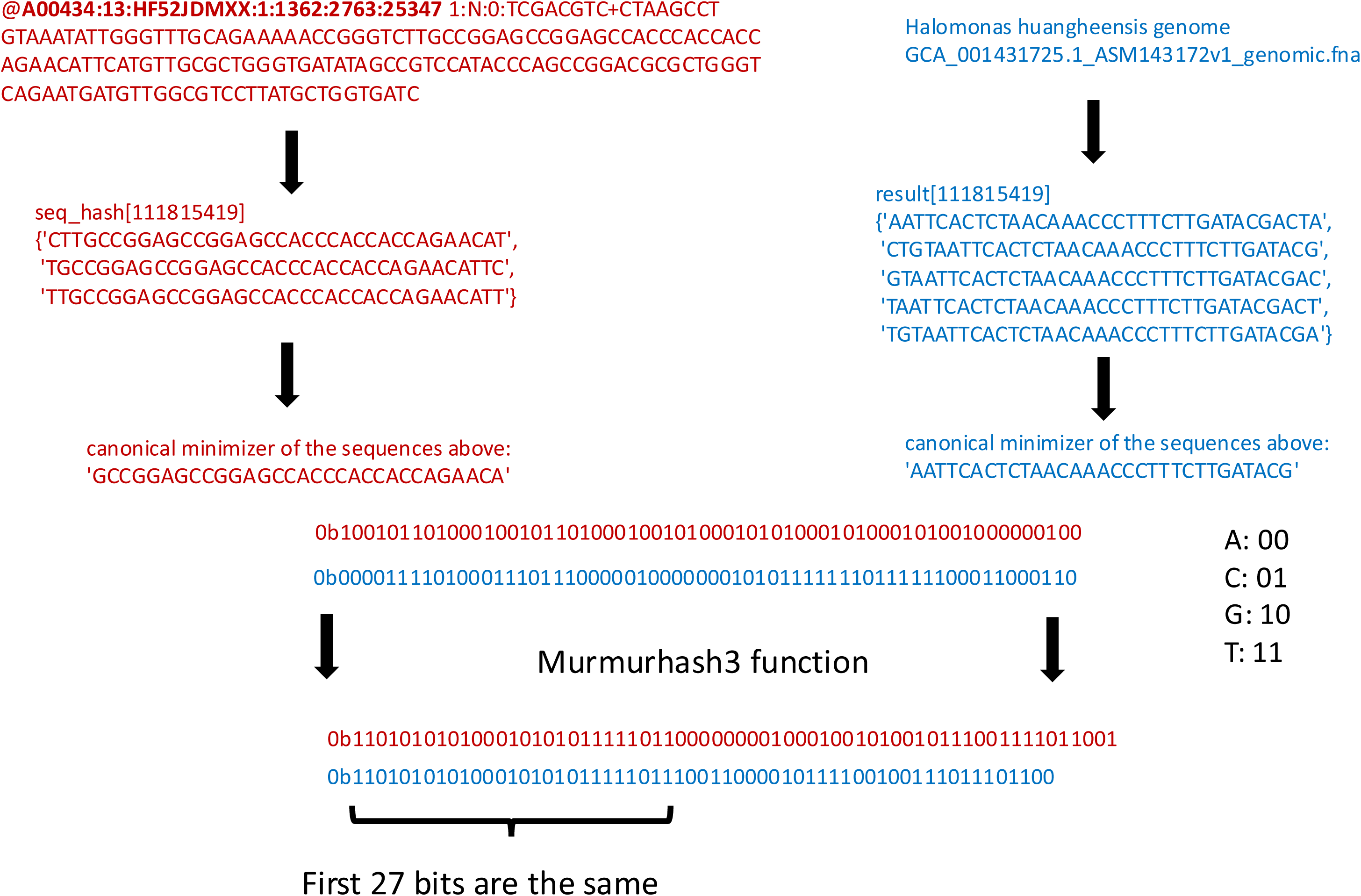
Example of misclassification from shared compact hash codes.

**Figure S2.**
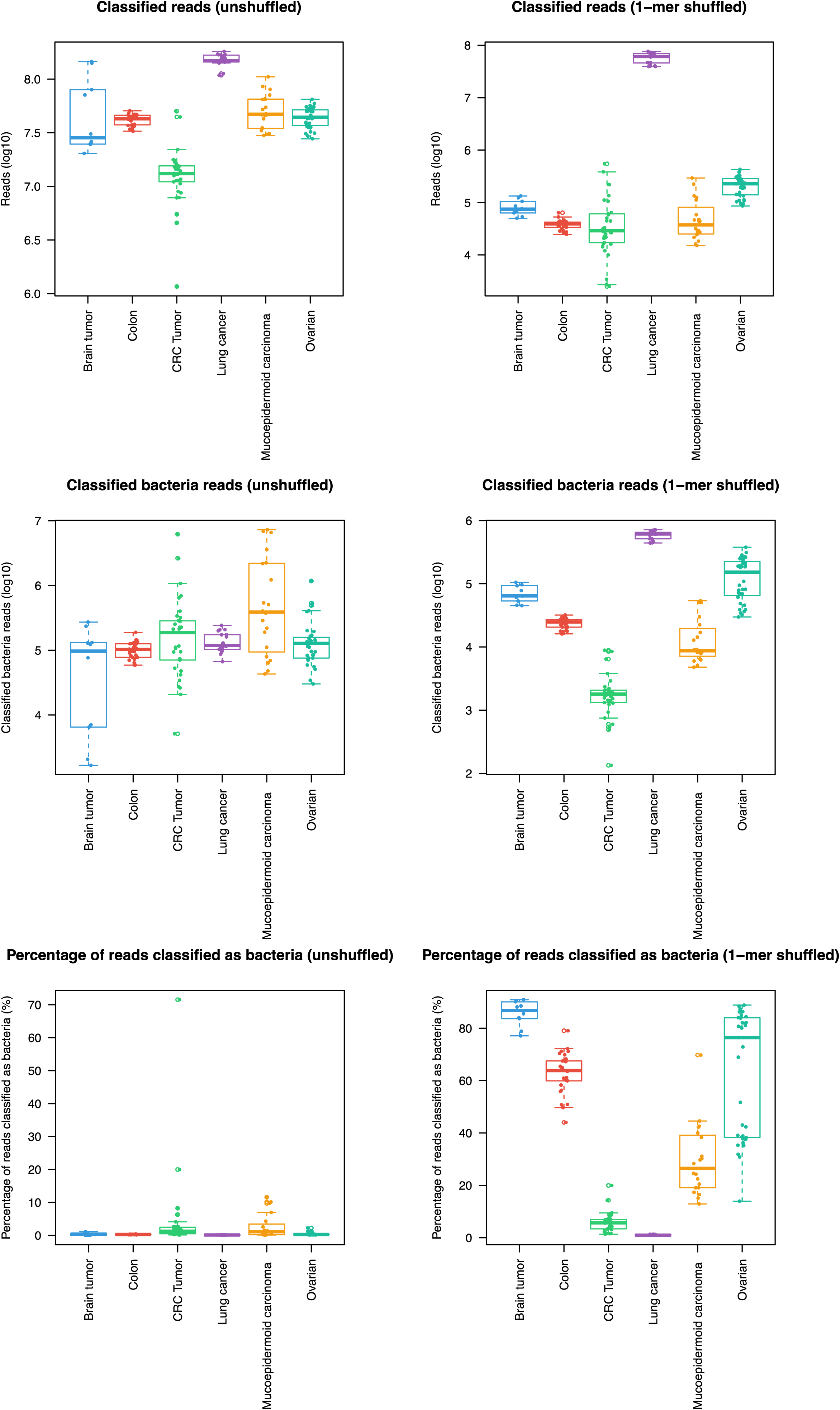
Classified total reads and bacteria reads by Kraken2.

**Figure S3.**
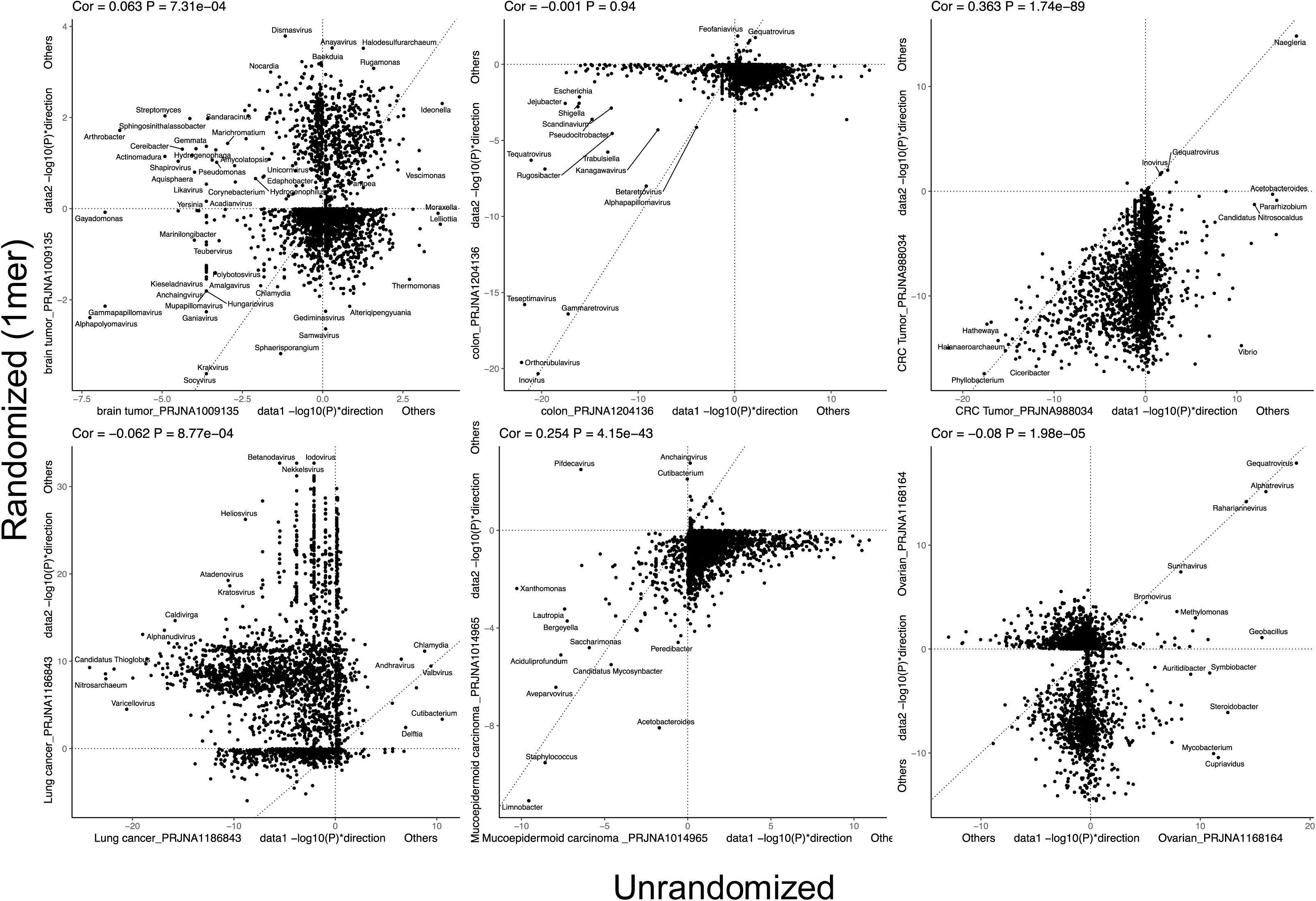
Comparison of statistical analyses results in randomized reads and unrandomized reads. Statistical analyses were performed to analyze the difference between one tumor type and the rest. X and Y axes are –log10(P-value) multiplied by +1/-1 to specify the direction of changes.

